# Daily HIV pre-exposure prophylaxis enhances monocyte activation and effector function and reprogrammes cellular metabolism

**DOI:** 10.64898/2026.01.08.698427

**Authors:** Grainne Jameson, Dearbhla M Murphy, Isabella Batten, Sarah A Connolly, Adam H. Dyer, Annmarie White, Sarah Cochrane, Dearbhla Murphy, Laura Quinn, Emma Devitt, Giovanni Villa, Sharee A Basdeo, Liam Townsend

**Affiliations:** School of Medicine, Trinity Translational Medicine Institute, St James’s Hospital, Trinity College Dublin, The University of Dublin, Ireland; Department of Genitourinary Medicine and Infectious Diseases, St James’s Hospital, Dublin, Ireland

## Abstract

**Background:** Pre-exposure prophylaxis (PrEP) with tenofovir/emtricitabine (TDF/FTC) is highly effective for HIV prevention. While antiretroviral therapy (ART) is linked to chronic inflammation in people living with HIV, its direct effects on immune phenotype, function, and metabolism in HIV-negative individuals remain unclear.

**Methods:** Gay, bisexual, and other men who have sex with men (gbMSM) on daily TDF/FTC PrEP underwent immunophenotyping and single-cell metabolic profiling using SCENITH™. Cytokine and chemokine responses were measured ex vivo and after lipopolysaccharide or *Mycobacterium tuberculosis* stimulation. These were compared with a demographically similar PrEP-naïve cohort, with five individuals followed 6–9 months after PrEP initiation.

**Findings:** Monocytes from people taking PrEP (n=15; median 533 days) exhibited higher activation marker expression (HLA-DR, CD14) *ex vivo* and enhanced IL-1β and TNF after bacterial challenge compared with PrEP-naïve individuals (n=11). Longitudinal follow-up indicated that PrEP initiation increased monocyte activation markers (HLA-DR, CD14, CD40, TNFRI/II) and cytokine production (IL-1β, TNF, GM-CSF, IFN-γ, Granzyme B, MIP-1α). Reduced glucose dependency was observed in monocytes, CD56^dim^ NK cells and CD4^+^ T cells 6–9 months after PrEP initiation.

**Interpretation:** Daily TDF/FTC promotes monocyte activation, enhances pro-inflammatory responses, and reprogrammes immune cell metabolism, highlighting ART’s potential to modulate immune-mediated inflammatory pathways in HIV-negative individuals.

**Funding:** *Trinity Translational Medicine Institute, Trinity College Dublin, Dublin, The Association of Physicians of Great Britain and Ireland, Health Research Board Ireland.*

**Graphical Abstract:** 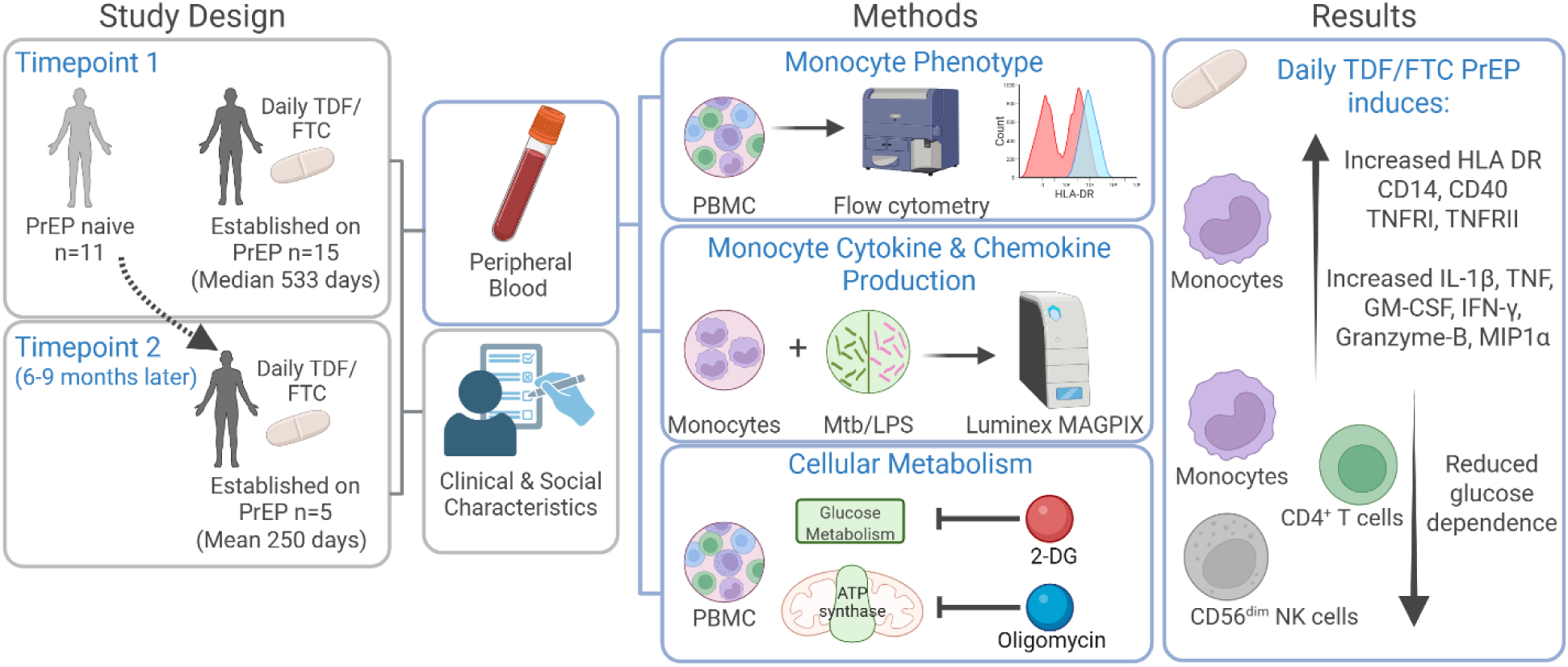

**Research in Context:** *Evidence before this study:* Despite effective viral suppression with antiretroviral therapy (ART), people living with HIV (PLWHIV) experience persistent immune activation and chronic inflammation that increase their risk of age-associated comorbidities. The extent to which ART itself contributes to this dysregulated immune state remains unclear, as HIV infection confounds most studies. Pre-exposure prophylaxis (PrEP) with tenofovir disoproxil fumarate and emtricitabine (TDF/FTC) offers a unique opportunity to examine the direct immunological effects of ART in HIV-negative individuals. However, few studies have explored how PrEP influences immune cell activation, effector function, or cellular metabolism in the absence of viral infection.

*Added value of this study:* This study is the first to characterise how daily PrEP with TDF/FTC modulates innate immune activation and cellular metabolism independently of HIV infection. We show that PrEP enhances monocyte activation and effector cytokine production while reprogramming metabolic pathways to reduce glucose dependency in monocytes, NK cells, and CD4⁺ T cells. These findings reveal previously unrecognised immunometabolic effects of ART in HIV-negative individuals, providing mechanistic insight into how ART exposure can shape immune homeostasis in the absence of infection.

*Implications of all the available evidence:* Our findings, together with existing data from PLWHIV, suggest that ART itself can remodel immune and metabolic pathways, potentially contributing to both beneficial and detrimental outcomes. ART-induced immunometabolic reprogramming may underlie persistent inflammation and the early onset of comorbidities observed in treated HIV infection, while also enhancing innate responsiveness to bacterial pathogens such as *Mycobacterium tuberculosis*. Understanding these mechanisms may inform future strategies to optimise ART regimens and mitigate inflammation-associated complications.

---

Keywords: ART, PrEP, immunometabolism, monocytes, HIV, tenofovir disoproxil fumarate

## Introduction

The evolution of therapies for Human Immunodeficiency Virus (HIV) infection has transformed the condition from a terminal illness to a chronic lifelong condition [1]. Lifespan for people living with HIV (PLWHIV) is now similar to their HIV-negative counterparts [2]. However, this increase in lifespan is not accompanied by a similar increase in healthspan, i.e. years lived in good health. PLWHIV develop ageing-related comorbidities at a younger age than HIV-negative individuals [3, 4]. The reasons behind this are multifactorial and incompletely understood. Some of the drivers of this accelerated ageing phenotype are co-infections such as viral hepatitis and lifestyle factors [5, 6]. Chronic inflammation due to HIV infection, even in the presence of effective anti-retroviral treatment (ART), has also been implicated in the development of ageing-associated comorbidities [7].

ART has been proposed to contribute to the immune dysregulation and altered metabolism that precedes the development of multimorbidity in PLWHIV [8]. The role of ART is particularly relevant due to its expanded use for prevention of HIV acquisition and allows us to determine the immunomodulatory effects of ART (in the absence of HIV) in otherwise healthy individuals. HIV pre-exposure prophylaxis (PrEP), usually as a fixed-dose combination of two nucleoside reverse transcriptase inhibitors (NRTIs) tenofovir disoproxil fumarate (TDF) and emtricitabine (FTC), taken either daily or on-demand, is highly effective at preventing HIV infection in individuals at increased risk of HIV acquisition [9, 10]. The majority of PrEP users are gay, bisexual and men who have sex with men (gbMSM). There are few studies examining the impact of PrEP on immune responses. Individuals taking PrEP do not develop enduring infection or antibody responses to HIV, despite presumed ongoing exposure to the virus. It is likely that the innate immune response is still activated in this context, and may be modulated by PrEP [11]. Additionally, TDF has been demonstrated to alter host cytokine and interferon responses [12, 13].

We hypothesised that PrEP alters innate immune activation and responses to unrelated bacterial stimuli mechanistically associated with changes in cellular metabolism. Previous *in vitro* studies have demonstrated altered monocyte phenotype and function when exposed to TDF/FTC [14, 15]. We compared immune responses in gbMSM taking PrEP to gbMSM with similar lifestyle risk factors who were not taking PrEP. To assess differences in immune responses, we examined monocyte, natural killer (NK) cell and T cell phenotype, function and metabolism in a cohort of gbMSM already established on daily PrEP (n=15) and a cohort of gbMSM prior to commencing PrEP (n=11). We then prospectively followed the cohort commencing PrEP and re-assessed these immune parameters 6-9 months after they were established on daily PrEP (n=5).

## Methods

### Study setting, participants and clinical covariates

All participants were recruited through the PrEP service at St James’s Hospital, Dublin, Ireland, which is a large tertiary referral centre. The inclusion criteria were cisgender gbMSM aged >18 years of age already established on daily PrEP or attending to commence daily PrEP. Exclusion criteria were any participant who self-reported symptoms of any infection, or detection of an asymptomatic sexually transmitted infection (STI) at time of recruitment. Participants taking event-based PrEP were excluded. Active injection drug use, receipt of any vaccine in the preceding three months, and receipt of immunomodulatory medication were also exclusion criteria. Sociodemographic data collected included age, ethnicity, number of sexual partners in the preceding month and preceding six months, condom usage, and STI history in the preceding twelve months. Substance misuse was recorded. Comorbidities and comedications were also recorded. All participants underwent screening for STIs, regardless of symptoms, with a first void urine, a rectal swab and pharyngeal swab for *Chlamydia trachomatis* and *Neisseria gonorrhoea* polymerase chain reaction testing. Serology for HIV, Hepatitis B, Hepatitis C, and *Treponema pallidum* was also performed. 40 ml of venous blood was collected in lithium heparin tubes. Participants commencing daily PrEP were invited to return for repeat sampling after a minimum of six months of daily PrEP, with identical sociodemographic and biological parameters recorded.

### Peripheral blood mononuclear cell and monocyte isolations

Peripheral blood mononuclear cells (PBMC) were isolated from whole blood samples by density centrifugation over Lymphoprep^TM^ (StemCell Technologies). Monocytes were separated using the EasySep^TM^ Human Monocyte Enrichment Kit without CD16 Depletion (19058; STEMCELL) according to the manufacturer’s instructions. Monocytes were added to a non-treated 48 well tissue culture plate (200,000 cells/well). Cells were stimulated with *Mycobacterium tuberculosis* (Mtb) whole cell lysate (10 μg/ml; BEI Resources) or lipopolysaccharide (LPS; 10 ng/ml; Sigma Aldrich) for 18 hours. Cells were left unstimulated as a control.

### Cytokine and chemokine analyses

A Human Luminex Discovery Kit (Cat LXSAHM-25) was custom designed to assess the following analytes in monocyte supernatants: TNF, IL-10, MCP-1, IL-1β, MIP-1α, IL-17α, GM-CSF, G-CSF, IFN-γ, and Granzyme B. The assay was carried out as per manufacturer’s protocol and analysed on the Luminex MAGPIX system. Unstimulated and stimulated cells from each donor at each timepoint were analysed concurrently on the same plate, thereby eliminating potential inter-assay variability. Analyte concentrations were determined based on standards supplied in the kit with a 5-parameter logistic regression curve used in Xponent^TM^ software. To minimise batch effects across the two time points, all stimulations were prepared prior to study initiation and thawed on the day of use to enhance consistency. To ensure maximal consistency and minimize technical variation, Luminex analyses of T1 and T2 supernatants were performed simultaneously using the same patient samples on the same assay plate.

### Immune cell phenotyping

Phenotypic analyses were carried out using flow cytometry. PBMC (100,000/tube) were resuspended in PBS (Gibco) and stained immediately. For lymphocyte phenotyping, cells were stained for 10 minutes at room temperature with anti-CD56 BV421 (1:100; 5.1H11), anti-CD69 BV510 (1:50; FN50), anti-CD8 BV650 (1;100; SK1), anti-CD27 BV785 (1:50;LG.3A10), anti-CD45RO APC-Cy7 (1:33;UCHL1), anti-CD49a FITC (1:50; TS2/7), anti-CXCR6 PerCP-cy5.5 (1:50; K041E5), anti-CD16 PE-Cy7 (1:200; 3G8), anti-NKG2C PE (1:100; S19005E), anti-CD3 PE-Dazzle (1:100; OKT3), and anti-NKG1A APC (1:100; S19004C) (all BioLegend).

For monocyte phenotyping, cells were stained for 10 minutes at room temperature with anti-CD86 BV421 (1:100;IT2.2), anti-CD45RA BV711 (1:100; HI100), anti-CD40 APC-Cy7 (1:33; HI100), anti-HLA-DR APC (1:100; L243), anti-CD14 AlexFluor488 (1:100; HCD14), anti-CD16 PerCP-Cy5.5 (1:100; 3G8), anti-CD80 PE-Cy7 (1:100; 2D10), anti-TNFRI PE (1:50; W15099A), and anti-TNFRII PE-Dazzle (1:100; 3G7A02) (all BioLegend). After staining, cells were washed, fixed and stored at 4°C until acquisition on the BD LSR Fortessa. Data was analysed using FlowJo^TM^ version 10.1.

### Measuring immune cell metabolism using SCENITH

The SCENITH protocol uses puromycin as a surrogate for protein synthesis as it becomes incorporated into newly formed proteins **(Figure 1)** [16, 17]. Immediately following isolation, SCENITH metabolic function profiling of PBMC was carried out. Cells (200,00/tube) were treated for 40 minutes at 37°C, 5% CO_2_ with DMSO control, 2-deoxy-D-glucose (2-DG 100 mM; Sigma-Aldrich), oligomycin (1 μM; Sigma-Aldrich), a combination of both drugs, or with Harringtonine (2 μg/ml; Sigma-Aldrich), which inhibits protein synthesis. Puromycin (10 μg/ml; Sigma-Aldrich) was added for the final 35 minutes. Cells were then washed with PBS and surface stained with anti-CD16 PE-Cy7 (1:200; 3G8), anti-CD56 BV421 (1:100; 5.1H11), anti-CD14 AlexFluor488 (1:100; HCD14), CD4 PerCP-Cy5.5 (1:100; RPA-T4), CD3 BV510 (1:100; UCHT1), Zombie-NIR (1:200) and

**Figure 1.**
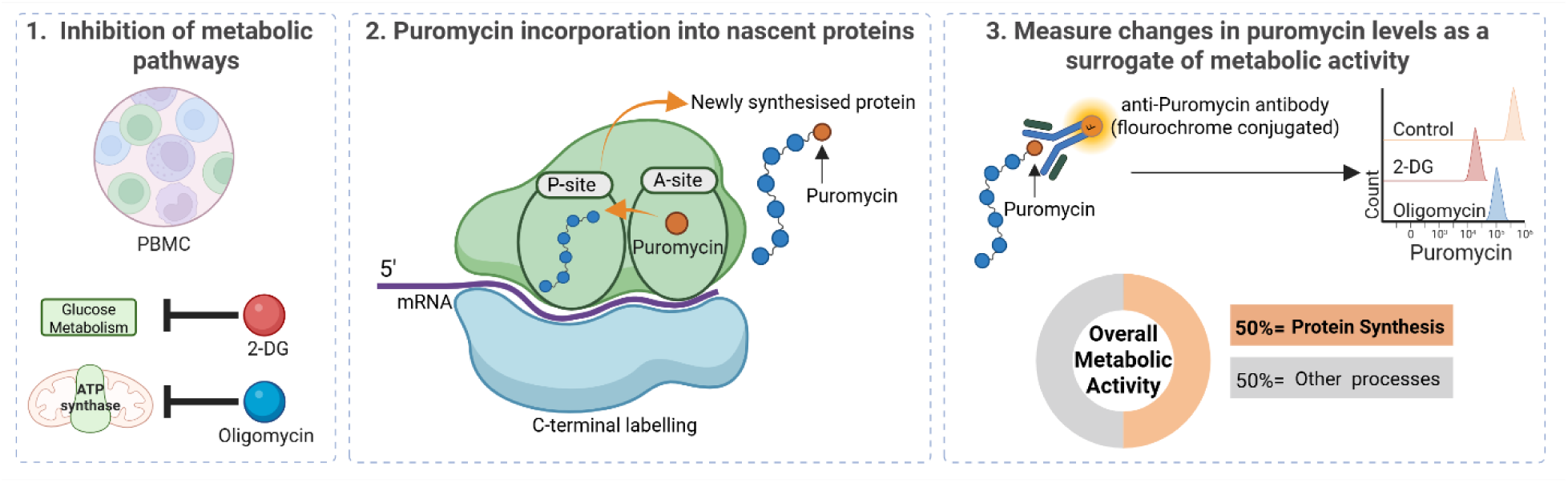
SCENITH protocol overview. Peripheral blood mononuclear cells (PBMC) are treated with metabolic inhibitors targeting specific pathways: 2-deoxyglucose (2-DG) to inhibit glucose metabolism and oligomycin to inhibit mitochondrial ATP synthase. Puromycin, an analogue of aminoacyl-tRNA, is incorporated into nascent polypeptide chains at the ribosomal A-site, resulting in C-terminal labelling of newly synthesized proteins. Incorporated puromycin is then detected by flow cytometry using a fluorochrome-conjugated anti-puromycin antibody, providing a quantitative measure of overall cellular metabolic activity. As protein synthesis accounts for approximately 50% of total metabolic activity, puromycin incorporation serves as a reliable surrogate for assessing cellular metabolic state.

Fc Block (1:100) (all Biolegend) for 10 minutes at room temperature. After staining, cells were washed with PBS and fixed using fixation/permeabilisation buffer from the FoxP3/Transcription Factor Staining Buffer Set (eBioscience) for 15 minutes in the dark at room temperature. The cells were then washed with permeabilisation buffer. Cells were stained intracellularly with anti-Puromycin (1:50; BioLegend) diluted in permeabilisation buffer with 20% foetal bovine serum and incubated in the dark at 4°C for 30 minutes. Cells were then washed once with permeabilisation buffer and resuspended in PBS for acquisition on the BD LSR Fortessa. Data was analysed using FlowJo^TM^ version 10.1.

Cellular dependency on mitochondrial respiration and glycolytic capacity was estimated by measuring puromycin incorporation post oligomycin treatment. Cellular dependence on glucose, fatty acid oxidation and amino acid oxidation was estimated by measuring puromycin incorporation post 2-DG treatment. Mitochondrial Dependence (MD) was calculated using the following equation: =100(Co-O)/(Co-DGO). Glycolytic Capacity (GC) was calculated using the following equation: =100-100(Co-O)/(Co-DGO).

### Statistical analyses

Cohort characteristic data was analysed using Stata version 19.0 (Stata Statistical Software). All other statistical analyses were performed using GraphPad Prism 10.4.2 software. Statistically significant differences between two normally distributed groups were determined using Student’s paired or unpaired t-tests with two-tailed P-values, unless a one-tailed statistical test is explicitly stated in the figure legend. Differences between two or more groups containing more than one variable were determined by two-way ANOVA with Sidak’s multiple comparisons test or Tukey’s multiple comparisons test. P-values of ≤0.05 were considered statistically significant and denoted using an asterisk. Details of the specific statistical test performed can be found in the figure legends.

### Role of the funding source

The funding sources had no involvement in the study design; in the collection, analysis, and interpretation of data; in the writing of the report; and in the decision to submit the paper for publication.

## Results

### Participant characteristics

There were n=26 participants recruited, with n=15 established on daily PrEP and n=11 recruited prior to commencing PrEP (treatment naive). All were gbMSM, with a mean age of 34.4 years. The only comorbidities recorded were asthma, depression and hyperlipidaemia. The only prescribed medications in the cohort were salbutamol, escitalopram and atorvastatin. Participants (n=11) reported at least one STI in the preceding 12 months; n=8 *N. gonorrhoea,* n=6 *C. trachomatis,* n=1 syphilis and n=1 L*ymphogranuloma venereum*. The demographics between the PrEP and non-PrEP cohorts were similar, with no differences in age, sexual behaviours, or substance misuse (**Table 1**). The median time on TDF/FTC for the cohort taking PrEP was 533 days (IQR 273 – 1,449).

**Table 1:**
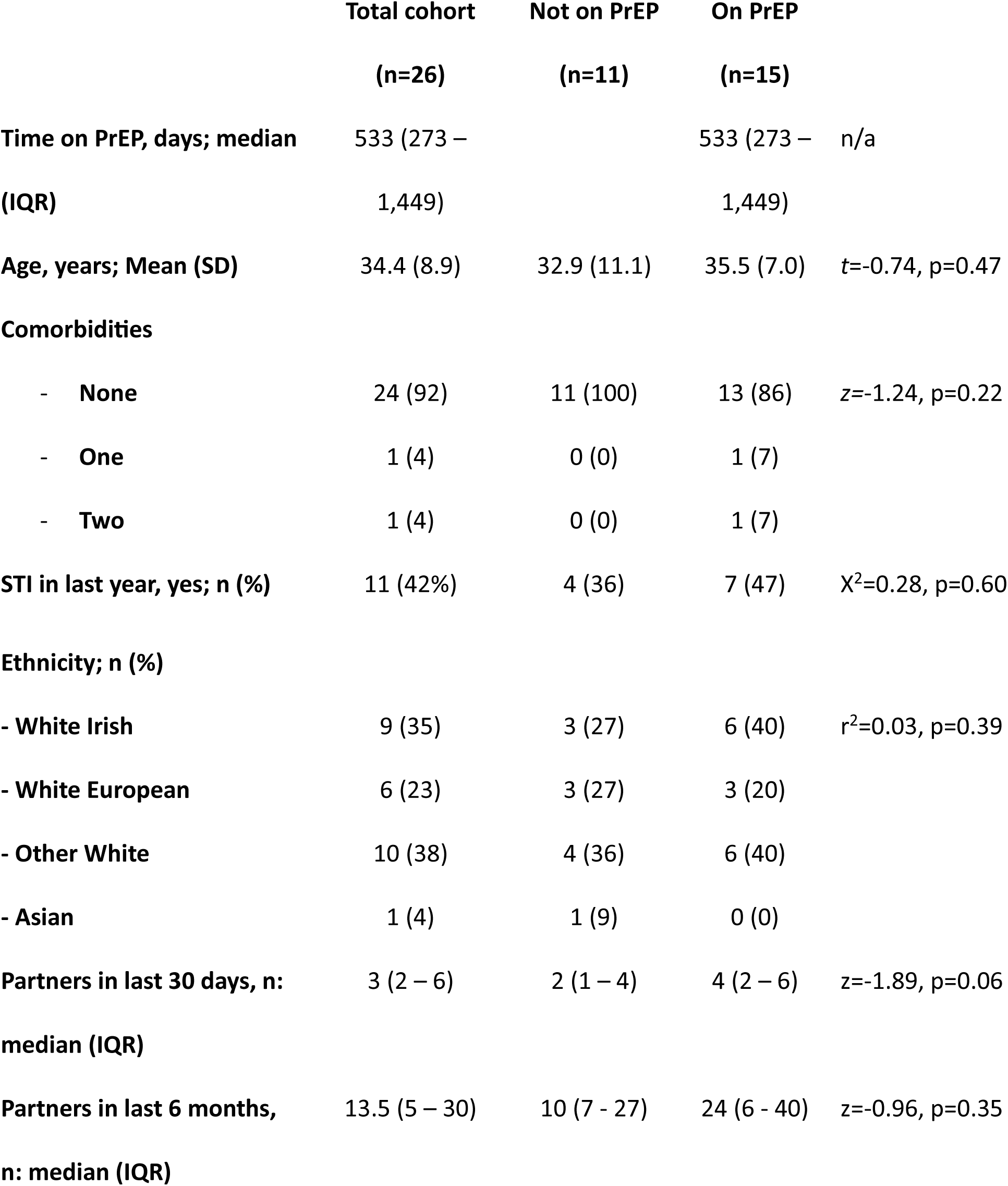
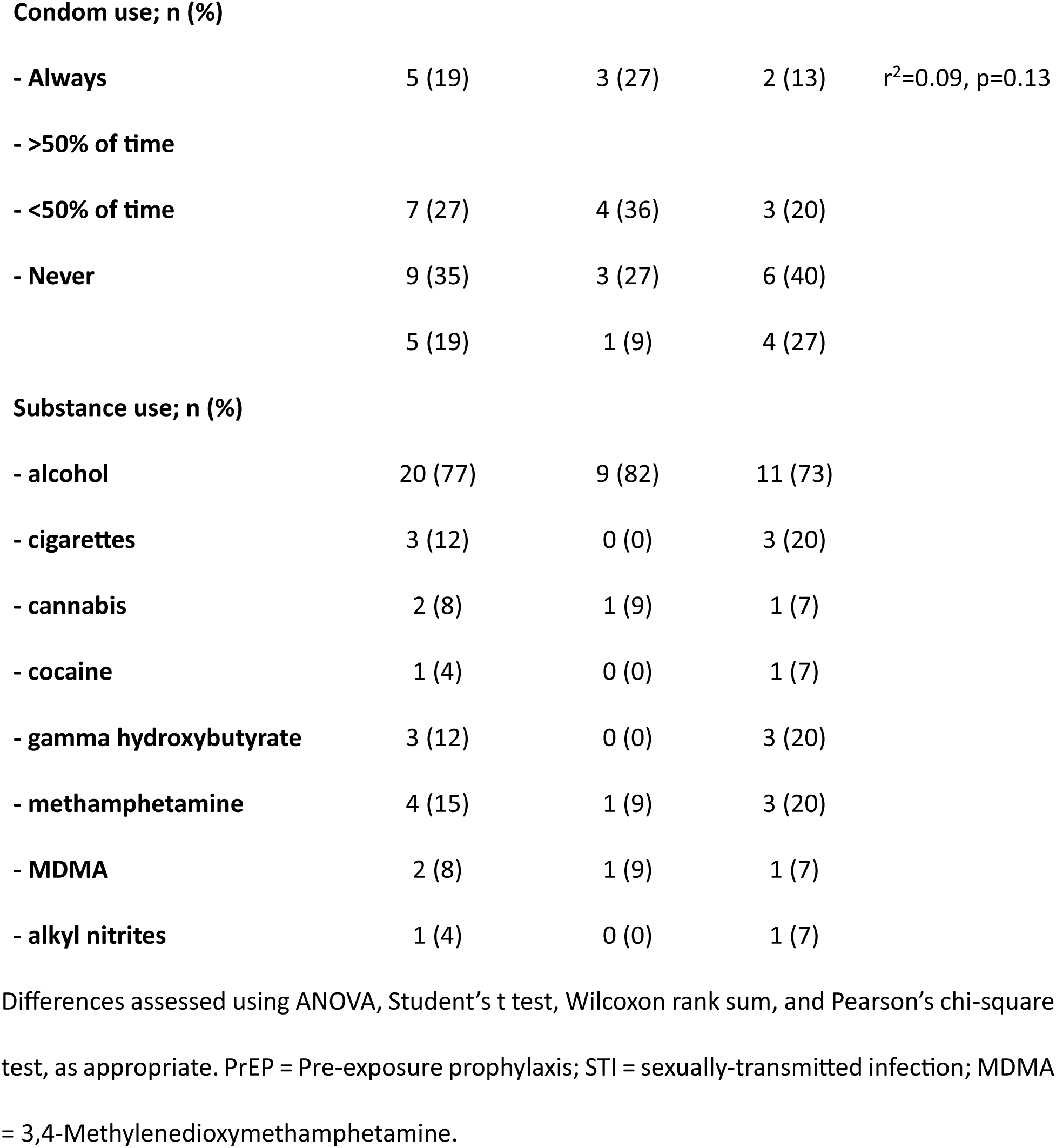
Cohort characteristics.

|  | **Total cohort** | **Not on PrEP** | **On PrEP** |  |
| --- | --- | --- | --- | --- |
|  | **(n=26)** | **(n=11)** | **(n=15)** |  |
| **Time on PrEP, days; median (IQR)** | 533 (273 – 1,449) |  | 533 (273 – 1,449) | n/a |
| **Age, years; Mean (SD)** | 34.4 (8.9) | 32.9 (11.1) | 35.5 (7.0) | $t=-0.74$, $p=0.47$ |
| **Comorbidities** |  |  |  |  |
| - **None** | 24 (92) | 11 (100) | 13 (86) | $z=-1.24$, $p=0.22$ |
| - **One** | 1 (4) | 0 (0) | 1 (7) |  |
| - **Two** | 1 (4) | 0 (0) | 1 (7) |  |
| **STI in last year, yes; n (%)** | 11 (42%) | 4 (36) | 7 (47) | $\chi^2=0.28$, $p=0.60$ |
| **Ethnicity; n (%)** |  |  |  |  |
| - **White Irish** | 9 (35) | 3 (27) | 6 (40) | $r^2=0.03$, $p=0.39$ |
| - **White European** | 6 (23) | 3 (27) | 3 (20) |  |
| - **Other White** | 10 (38) | 4 (36) | 6 (40) |  |
| - **Asian** | 1 (4) | 1 (9) | 0 (0) |  |
| **Partners in last 30 days, n: median (IQR)** | 3 (2 – 6) | 2 (1 – 4) | 4 (2 – 6) | $z=-1.89$, $p=0.06$ |
| **Partners in last 6 months, n: median (IQR)** | 13.5 (5 – 30) | 10 (7 - 27) | 24 (6 - 40) | $z=-0.96$, $p=0.35$ |
| **Condom use; n (%)** |  |  |  |  |
| - Always | 5 (19) | 3 (27) | 2 (13) | $r^2=0.09$, $p=0.13$ |
| - >50% of time |  |  |  |  |
| - <50% of time | 7 (27) | 4 (36) | 3 (20) |  |
| - Never | 9 (35) | 3 (27) | 6 (40) |  |
|  | 5 (19) | 1 (9) | 4 (27) |  |

**Substance use; n (%)**
|  |  |  |  |
| --- | --- | --- | --- |
| - alcohol | 20 (77) | 9 (82) | 11 (73) |
| - cigarettes | 3 (12) | 0 (0) | 3 (20) |
| - cannabis | 2 (8) | 1 (9) | 1 (7) |
| - cocaine | 1 (4) | 0 (0) | 1 (7) |
| - gamma hydroxybutyrate | 3 (12) | 0 (0) | 3 (20) |
| - methamphetamine | 4 (15) | 1 (9) | 3 (20) |
| - MDMA | 2 (8) | 1 (9) | 1 (7) |
| - alkyl nitrites | 1 (4) | 0 (0) | 1 (7) |

Differences assessed using ANOVA, Student's t test, Wilcoxon rank sum, and Pearson's chi-square test, as appropriate. PrEP = Pre-exposure prophylaxis; STI = sexually-transmitted infection; MDMA = 3,4-Methylenedioxymethamphetamine.

### PrEP significantly enhanced monocyte proinflammatory phenotype and function

Cellular frequencies and phenotype of immune cell subsets present in PBMC from PrEP and non-PrEP participants were measured by flow cytometry. The frequencies of monocytes and lymphocytes were comparable between the two groups **(Figure 2A-B).** Similarly, no significant differences were observed in the frequencies of NK and T cells or their respective subsets, including CD56^bright^, and CD56^dim^ NK cells, and CD4^+^ and CD8^+^ T cells **(Figure 2C-E).** However, PrEP was associated with a pro-inflammatory phenotype in peripheral blood monocytes, characterised by increased expression of the antigen-presentation molecule HLA-DR and the TLR-4 co-receptor CD14 (**Figure 2F – H**). There were no differences observed between groups in monocyte expression of costimulatory molecules CD40, CD45RA, CD80 and CD86, or TNF receptors; TNFRI and TNFRII (**Figure 2I – L**). Extensive phenotyping of NK and T cell activation markers was also performed however no significant differences were observed (**Supplemental Figure 1A-D**).

**Figure 2.**
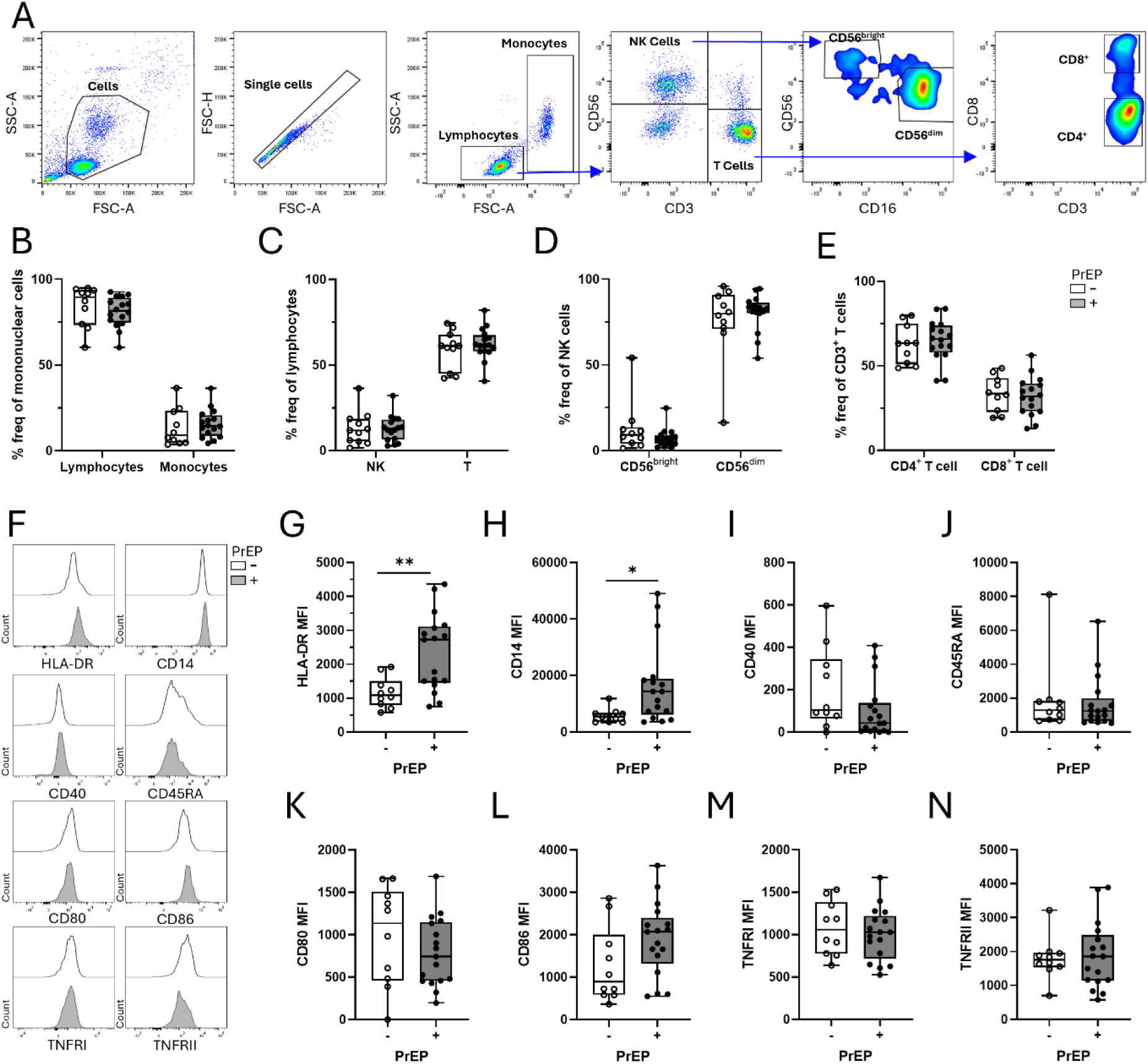
Daily PrEP induces a pro-inflammatory phenotype in peripheral blood monocytes. PBMC were isolated from peripheral blood of PrEP naive individuals (white; n=11) or individuals established on PrEP (grey; n=15) and stained with fluorochrome conjugated antibodies for CD14, CD56, CD3, CD4, CD8, HLA-DR, CD40, CD45RA, CD80, CD86, TNFRII and TNFRII for flow cytometric analysis. **(A)** Representative flow plot of gating strategy. **(B)** Lymphocyte and monocyte frequencies as a percentage of total mononuclear cells. **(C)** NK and T cell frequencies as a percentage of total lymphocytes. **(D)** CD56^bright^ and CD56^dim^ frequencies as a percentage of total NK cells. **(E)** CD4^+^ and CD8^+^ frequencies as a percentage of total T cells. **(F)** Representative histograms of the MFI of the activation markers on monocytes. Expression levels of **(G)** HLA-DR, **(H)** CD14, **(I)** CD40, **(J)** CD45RA, **(K)** CD80, **(L)** CD86, **(M)** TNFRI or **(N)** TNFRII on total monocytes. Each dot represents an individual donor, and data were analysed using (B-E) a two-way ANOVA with Šidák’s multiple comparisons test or (G-N) an unpaired t test where *P<0.05, or **P<0.01.

Given the changes in monocyte markers of activation, monocyte function was next investigated. Monocytes were isolated from PBMC and left unstimulated or stimulated with Mtb lysate or LPS for 18 hours. Production of IL-1β, TNF, GM-CSF, MIP-1α, MCP-1, Granzyme-B, IFN-γ, IL-17, IL-10 and G-CSF were measured under basal conditions (unstimulated), following TLR4 activation by LPS, and following multiple PRR activation with Mtb lysate. Under basal conditions, there were no changes in monocyte cytokine or chemokine production between groups (**Figure 3A, Supplemental Figure 2A**). Following stimulation of TLR4 with LPS, increased cytokine and chemokine production was observed. The magnitude of these responses were similar in both the PrEP and non-PrEP groups (**Figure 3B, Supplemental Figure 2B**). However, following simultaneous stimulation of multiple PRRs with Mtb lysate, monocytes in the PrEP cohort demonstrated significantly higher production of IL-1β and TNFα compared with -PrEP naive participants (**Figure 3C**). GM-CSF, MIP-α and MCP-1 were not signficantly altered in individuals initiated on PrEP compare with naive controls (Figure 3C). This suggests that daily PrEP results in enhanced production of pro-inflammatory IL-1β and TNFα in circulating monocytes. Cytokine and chemokine production following Mtb stimulation, including Granzyme B, IFN-γ, IL-17, IL-10, and G-CSF, was extensively profiled but showed no significant changes. (**Supplemental Figure 2C)**.

**Figure 3.**
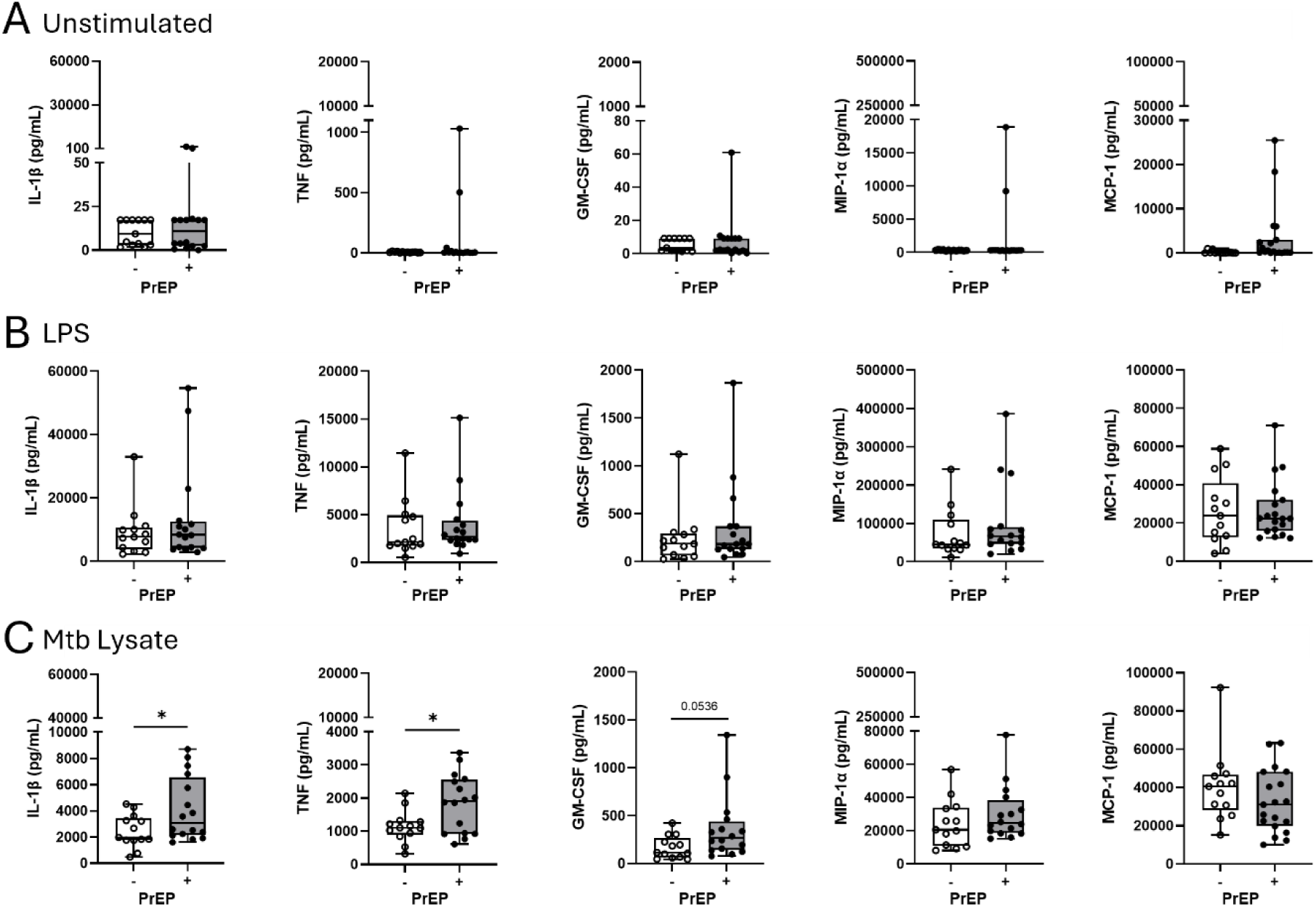
Daily PrEP enhances monocyte pro-inflammatory responses to Mtb lysate. Monocytes were magnetically sorted from the PBMC of PrEP naive individuals (white; n=11) or individuals established on PrEP (grey; n=15) and **(A)** left unstimulated, **(B)** stimulated with LPS or **(C)** stimulated with Mtb lysate for 18 hours. The concentrations (pg/ml) of IL-1β, TNF, GM-CSF, MIP-1α or MCP-1 in the supernatant were measured using a Luminex Discovery assay. Each dot represents an individual donor, and data were analysed using an unpaired t test where *P<0.05.

### Longitudinal analysis following treatment initiation indicates that PrEP significantly increased monocyte pro-inflammatory phenotype and function

Due to the differences in monocyte phenotype and effector function observed between PrEP and non-PrEP participants we next prospectively followed the non-PrEP participants who commenced daily PrEP post timepoint 1. These n=11 participants were invited to return for repeat sampling a minimum of six months after PrEP commencement. Six participants were lost from the study; n=2 had switched to event-based dosing, n=3 declined take part/had disengaged from the service, and n=1 had a positive STI screen at Timepoint 2, leaving n=5 eligible participants. These patients had been taking PrEP for a mean of 250 days (SD 10.8). There were no demographic differences between the eligible n=5 and the six participants who did not provide a timepoint 2 sample.

Once established on daily PrEP, markers of monocyte activation, namely HLA-DR, CD14, CD40, TNFRI and TNFRII expression levels were significantly higher compared with timepoint 1 (**Figure 4A**). Following Mtb stimulation, there were marked increases across a range of cytokines produced by monocytes, with significantly higher concentrations of IL-1β, TNF, GM-CSF, IFN-γ, Granzyme-B, and MIP-1α, produced compared with their own matched PrEP-naïve control from timepoint 1 (**Figure 4B, Supplemental Figure 3C**). Monocyte cytokine and chemokine production were not significantly altered post initiation of PrEP under basal conditions, or after stimulation with LPS (**Supplemental Figure 3A-B**).

**Figure 4.**
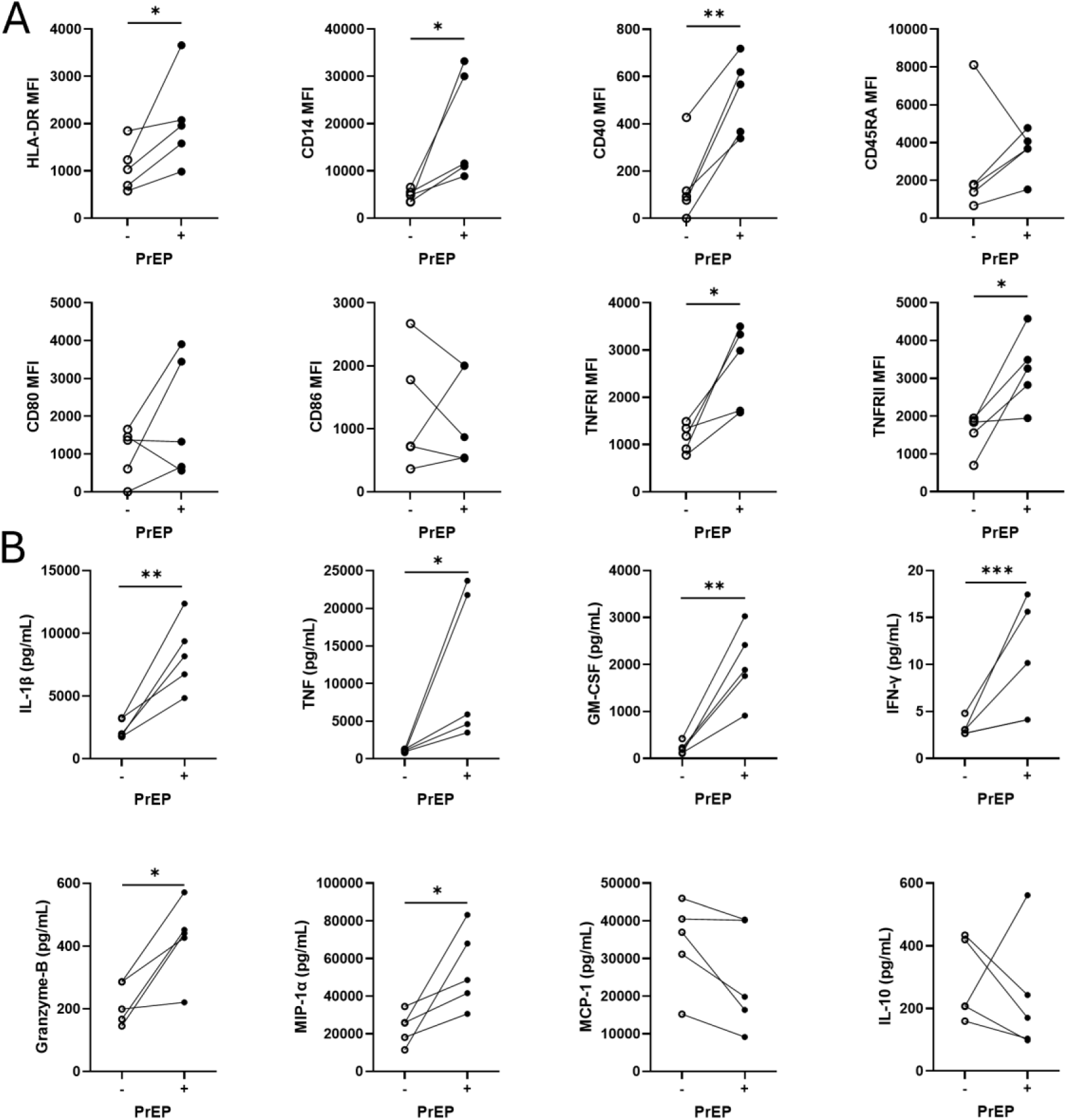
Longitudinal follow up confirms PrEP-treatment effects. **(A)** PBMC were isolated from the matched blood (n=5) of PrEP naïve individuals at T1 (-; open circle) who were established on PrEP at T2 (+; closed circle) and stained with fluorochrome conjugated antibodies for CD14, HLA-DR, CD40, CD45RA, CD80, CD86, TNFRII and TNFRII for flow cytometric analysis. **(A)** Expression levels of HLA-DR, CD14, CD40, CD45RA, CD80, CD86, TNFRI or TNFRII on total monocytes. **(B)** Magnetically-sorted monocytes were stimulated with Mtb lysate for 18 hours. The concentrations (pg/ml) of IL-1β, TNF, GM-CSF, IFN-γ, Granzyme-B, MIP-1α, MCP-1, and IL-10 in the supernatant were measured using a Luminex Discovery assay. Each dot represents matched data at each timepoint joined by a line. Data were analysed using a paired t test where *P<0.05, **P<0.01, or ***P<0.001. For the HLA-DR, CD14 and TNF statistical analyses, a one-tailed paired t test was used.

### PrEP metabolically reprogrammes myeloid and lymphoid cell compartments

Since innate immune function is integrally linked to cellular metabolism, we hypothesised that the observed phenotypic and functional changes in monocytes from people established on PrEP may be a result of a PrEP-induced metabolic reprogramming. Therefore, we next investigated whether PrEP alters monocyte metabolic profiles using the SCENITH metabolism by flow cytometry protocol [16] (**Figure 1**). We took advantage of this protocol’s ability to assess metabolic profiles at a single cell resolution and analysed CD56^bright^ NK cells, CD56^dim^ NK cells, CD4^+^ T cells and CD8^+^ T cells. Using puromycin incorporation as a surrogate marker of metabolic activity, we assessed overall cellular metabolism in PrEP-naïve individuals and again 6–9 months after PrEP initiation. No significant changes in overall metabolic activity were observed across immune cell subsets between timepoints **(Figure 5A-B).** To evaluate glucose dependency, cells were treated with the glucose analogue 2-deoxyglucose (2-DG) to inhibit glucose metabolism. PrEP initiation resulted in a reduction in glucose dependency in monocytes, CD56^dim^ NK cells and CD4^+^ T cells, but remained unchanged in CD56^bright^ NK cells and CD8^+^ T cells **(Figure 5C)**. Inhibition of mitochondrial respiration using oligomycin enabled assessment of mitochondrial dependency. These analyses revealed that PrEP establishment reduced mitochondrial dependency in CD56^bright^ NK cells while no significant changes were observed in monocytes, CD56^dim^ NK, CD4^+^, or CD8^+^ T cells **(Figure 5D)**.

**Figure 5.**
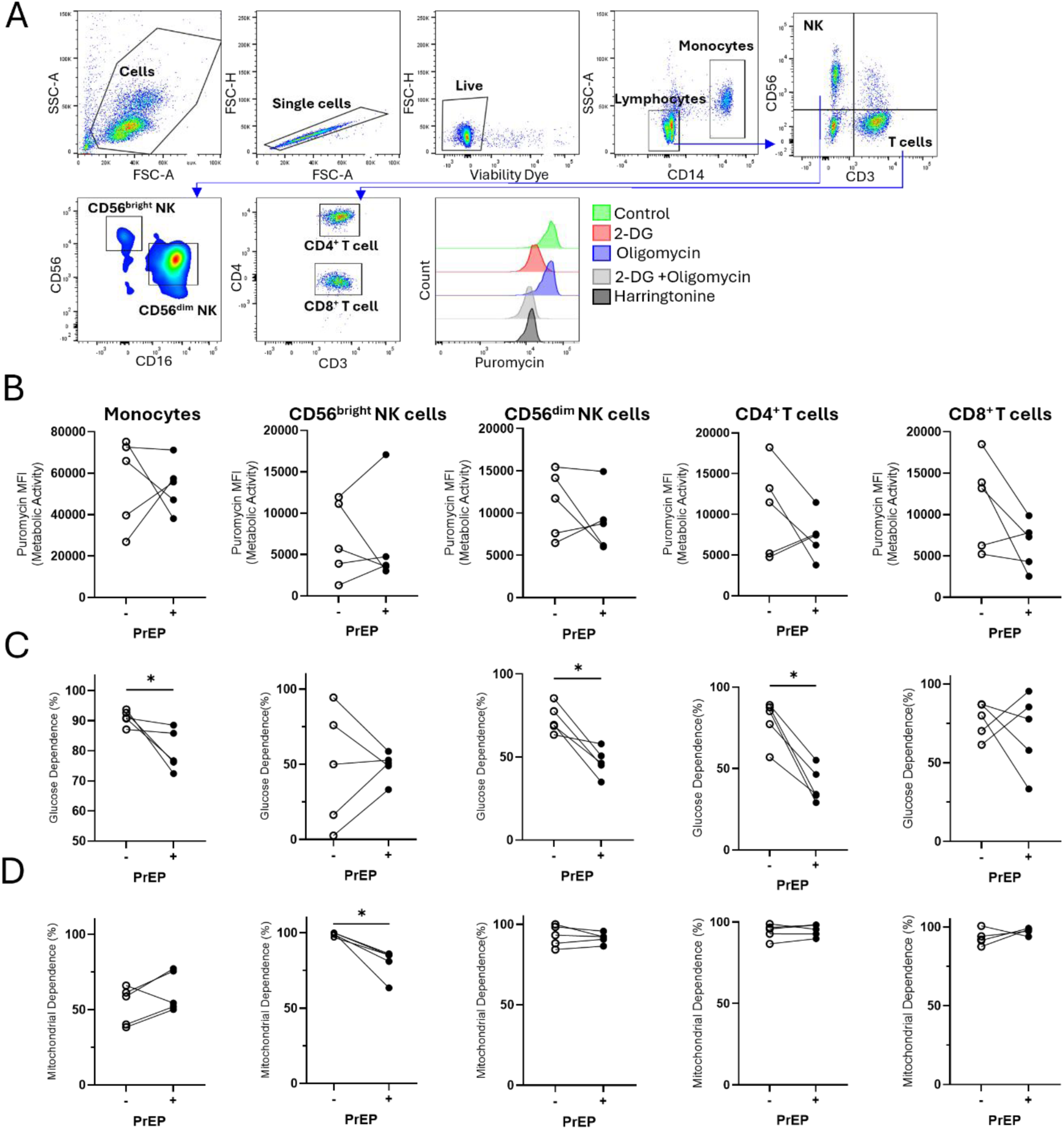
Daily PrEP induces metabolic reprogramming in myeloid and lymphoid cells. PBMC from (n=5) individuals not on PrEP at T1 (-; open circle) and on PrEP on T2 (+; closed circle) were treated with DMSO control or metabolic inhibitors 2-deoxyglucose (2DG), oligomycin (Oligo), or combinations of both, or harringtonine, which inhibits protein synthesis and acts as a negative control for this assay. Puromycin was then added, and cells were cultured for 40 min. Cells were washed and stained with fluorochrome-conjugated antibodies against CD14, CD56, CD3, CD4, CD8 and puromycin and analysed by flow cytometry. Analysis was performed on monocytes, CD56^bright^ NK cells, CD56^dim^ NK cells, CD4^+^ T cells and CD8^+^ T cells **(A)** Representative flow plot of gating strategy showing the analyses of monocytes and their puromycin staining across metabolic treatments. **(B)** Puromycin MFI values measuring overall metabolic activity in cells *ex vivo* across Ctrl (DMSO control), 2DG, and oligomycin treatments. **(C)** Percentage glucose dependence in cells *ex vivo*. **(D)** Percentage mitochondrial dependence in cells *ex vivo*. Each dot represents an individual donor and matched data at each timepoint joined by a line. Data were analysed using a paired t test where *P<0.05. For the CD56^dim^ NK cell and CD4^+^ T cell statistical analyses, a one-tailed paired t test was used.

Taken together, these data indicate that initiation of daily PrEP is associated with altered innate immune phenotype and function and changes in the metabolism of circulating myeloid and lymphoid cells.

## Discussion

The contribution made by HIV ART, specifically TDF and FTC, to altered inflammatory responses remain poorly understood. In this study, we demonstrate that daily TDF/FTC PrEP induces measurable changes in immune cell phenotype, function, and metabolism in otherwise healthy gbMSM, independent of HIV infection. Our findings indicate that PrEP not only exerts antiviral effects but may also modulate innate and adaptive immune responses *in vivo*, supporting previous in vitro studies that have highlighted the immunomodulatory properties of TDF/FTC [13, 18].

We observed that monocytes from individuals taking PrEP exhibit an activated, pro-inflammatory phenotype, characterised by increased expression of HLA-DR, CD14, CD40, TNFRI, and TNFRII. HLA-DR is a major histocompatibility complex class II molecule that mediates antigen presentation to CD4^+^ T cells, and its upregulation suggests enhanced capacity for antigen presentation and initiation of adaptive immune responses. CD14 and CD40 are critical for pathogen recognition and co-stimulation, respectively, indicating that monocytes in PrEP users may be primed for more efficient microbial sensing and T cell activation. Notably, increased CD40 expression has also been linked to enhanced monocyte adhesion and cardiovascular disease [19, 20]. TNFRI and TNFRII are receptors for TNF with complementary functions: TNFRI mediates classical pro-inflammatory signalling, whereas TNFRII has been implicated in immune regulation and survival signalling. Upregulation of these receptors suggests that monocytes are primed for robust TNF-mediated responses, potentially amplifying responsiveness to microbial stimuli while maintaining regulatory capacity. Enhanced TNFRI signalling in particular has been associated with persistent pro-inflammatory states [21]. Together, these findings provide insight into the impact of TDF/FTC on inducing a pro-inflammatory state in the peripheral myeloid compartment, consistent with observations of elevated plasma markers of monocyte activation in PLWHIV with undetectable viral loads and evidence of TDF/FTC-associated immune activation within the central nervous system [22, 23].

Functionally, monocytes from individuals taking PrEP produce higher levels of pro-inflammatory cytokines and chemokines, including IL-1β, TNF, GM-CSF, IFN-γ, Granzyme-B, and MIP-1α, following stimulation with Mtb lysate. Both IL-1β and TNF are key mediators of host defence against *M. tuberculosis*, promoting leukocyte recruitment and macrophage activation [24]; however, excessive production can contribute to immunopathology and tissue damage [25]. GM-CSF stimulates myeloid cell proliferation and enhances neutrophil recruitment, and IFN-γ activates macrophages to improve intracellular pathogen killing. Granzyme-B mediates cytotoxicity against infected cells, including Mtb-infected targets, whereas MIP-1α recruits monocytes and other immune cells to sites of infection, propagating inflammation [26–28]. These findings are in line with prior studies showing that nucleoside reverse transcriptase inhibitors can influence monocyte cytokine production and activation marker expression [14].

Basal cytokine levels in unstimulated monocytes remained largely unchanged, suggesting that PrEP may prime immune cells without inducing overt systemic inflammation under resting conditions. This selective enhancement of responsiveness may be beneficial for host defence, particularly against opportunistic infections such as Mtb, although elevated levels of pro-inflammatory cytokines, especially TNF and IL-1β, have been linked to chronic inflammation, accelerated cardiovascular disease, and worse outcomes in acute infections [29–31]. Tuberculosis remains the leading cause of death in PLWHIV globally, and initiation of ART in TB patients can trigger TB-associated immune reconstitution inflammatory syndrome (TB-IRIS). Transcriptomic analyses reveal that innate immune mediators, including TLR and IL-1 pathways, are strongly upregulated at the characteristic time of TB-IRIS onset (approximately 2 weeks post ART initiation) [32, 33]. Consistently, in our study, cytokine responses to LPS were stable, whereas Mtb lysate elicited markedly enhanced cytokine and chemokine production post-PrEP initiation. These data suggest that PrEP selectively primes monocytes for a more robust pro-inflammatory response to complex pathogen-associated molecular patterns (PAMPs), potentially enhancing host defence, but also raising the possibility of maladaptive inflammation in certain contexts [34].

Metabolic profiling using SCENITH revealed that PrEP initiation is associated with reduced glucose dependency in monocytes, CD56^dim^ NK cells, and CD4^+^ T cells, as well as reduced mitochondrial dependency in CD56^bright^ NK cells. This reduced glucose dependency likely reflects lower basal glycolytic activity, which may indicate an enhanced capacity to upregulate glycolysis in response to infection [35–37]. Consequently, the observed decrease in glucose reliance in unstimulated cells does not necessarily signify impaired innate immunity, but rather suggests that PrEP reprogrammes metabolic function and further studies are needed to evaluate the metabolic flexibility of these immune cells following challenge [38]. These findings indicate that TDF/FTC induces metabolic reprogramming across both myeloid and lymphoid compartments.

Interestingly, older NRTIs have well-recognised mitochondrial toxicities, and both TDF and FTC have been shown to induce mitochondrial dysfunction [39, 40]. Previous *ex vivo* work has demonstrated that TDF/FTC can modulate monocyte-derived macrophage metabolism, with reduced mitochondrial mass and increased reactive oxygen species production [41]. In the context of our study, these TDF/FTC-induced metabolic changes, together with a pro-inflammatory monocyte phenotype, may contribute to the low-grade chronic inflammation observed in PLWHIV. At the same time, our findings suggest that PrEP may enhance metabolic flexibility, potentially enabling immune cells to mount more robust responses to environmental stress or infection. Whether these metabolic shifts confer altered immune surveillance, enhanced protection against pathogens, or increased risk of maladaptive inflammation remains to be determined.

Our study has limitations. Understanding the aetiology of altered inflammation in PLWHIV is challenging due to the complex characteristics of patients, who often present with diverse social backgrounds, multiple comorbidities, prior exposure to potentially toxic ART, and health behaviours that may promote inflammation. We mitigated many of these confounders by carefully matching our PrEP and non-PrEP cohorts. The shared lifestyle risk factors between these groups allow control of sociodemographic influences [42, 43], and TDF/FTC are the only ART regimen these participants have been exposed to. Additionally, the mean age of 34.4 years reduces the potential impact of immunosenescence and the pro-inflammatory senescence-associated secretory phenotype [44].

The cohort was restricted to healthy gbMSM, and findings may not extrapolate to other populations, such as cisgender women or individuals with comorbidities. Larger studies, including women and powered to determine sex-specific effects of ART on immunity, are therefore warranted. We only included participants taking daily TDF/FTC as PrEP, so it remains unknown whether similar immune changes occur with event-based dosing or with alternative regimens, such as tenofovir alafenamide/emtricitabine or lenacapavir. We did not directly assess responses to viral challenges or HIV peptides *ex vivo*, nor did we determine cytomegalovirus serostatus, which could be a hidden confounder in immune response analyses. The impact of this is partially mitigated by the use of matched samples from the same participants over a relatively short (six to nine month) period. Finally, while our data demonstrate ongoing immune modulation associated with PrEP, the study was not powered to detect clinically significant events resulting from altered inflammatory responses. Collectively, these findings support the need for longitudinal PrEP registries to assess long-term outcomes and refine risk–benefit evaluations.

We demonstrate a shift in inflammatory responses associated with daily TDF/FTC pre-exposure prophylaxis for HIV. The durability of these changes, and whether they may contribute to the development of comorbidities linked to chronic inflammation, represents an important avenue for future study. Our comparison of individuals established on PrEP for a mean of 533 days versus those on PrEP for a mean of 250 days suggests that some inflammatory signatures are maintained over time, such as increased expression of CD14 and HLA-DR on circulating monocytes and enhanced IL-1β and TNF production following bacterial challenge. Conversely, CD40 expression appeared to be elevated at earlier timepoints (up to ∼250 days) but normalized with longer-term PrEP, as no difference was evident in those established on PrEP for ∼533 days compared with PrEP-naïve controls. These findings indicate that the effects of PrEP on monocyte phenotype and function may evolve over time, underscoring the need for longitudinal studies to better understand potential sub-cohorts at risk of pathological inflammatory responses.

In summary, our study demonstrates that daily TDF/FTC PrEP is associated with a pro-inflammatory monocyte phenotype, enhanced cytokine and chemokine responses to complex microbial stimuli, and metabolic reprogramming across myeloid and lymphoid compartments. These findings highlight previously unappreciated off-target immunomodulatory effects of PrEP in otherwise healthy individuals. While the long-term clinical implications of these changes remain to be determined, our data suggest that PrEP may both enhance innate host defence against pathogens and potentially contribute to low-grade inflammation over time. Importantly, the observed time-dependent effects on specific markers such as CD40 underscore the dynamic nature of immune modulation with ongoing PrEP exposure. Future longitudinal studies across diverse populations will be essential to clarify the durability, functional consequences, and clinical relevance of these immunometabolic shifts, ultimately informing the long-term risk–benefit profile of HIV PrEP.

## Contributors

*S.A.B. and L.T. conceived the study and hypotheses. G.J., D.M.M., I.B., S.A.C., A.W, S.C, D.M, L.Q., E.D., G.V., S.A.B and L.T. participated in data collection. L.T. and S.A.B. provided domain expert advice and evaluation of the project. G.J., D.M., I.B. and S.A.B. performed data and statistical analyses. All authors interpreted the data. G.J., D.M., S.A.B. and L.T. wrote the initial draft of the manuscript. All authors provided substantive critical reviews and approved of the submitted manuscript*.

## Conflict of Interests

The authors declare that they have no competing interests.

## Funding Source

*This work was supported by a Building Engagement in Health Research grant from Trinity Translational Medicine Institute, Trinity College Dublin, Dublin and an Emerging Clinical Researcher award from the Association of Physicians of Great Britain and Ireland. Health Research Board EIA-2019-010, Health Research Board ILP-POR-2022-033, Health Research Board ILP-2024-044.*

## Ethical Approval statement

Ethical approval for this study was obtained from the joint ethics committee of Tallaght University Hospital and St James’s Hospital (reference 2024 – Apr – 34633463). Written informed consent was obtained from all participants.

## Data Sharing Statement

Data will be made available from the corresponding author upon request.

---

**Supplemental Figure 1.**
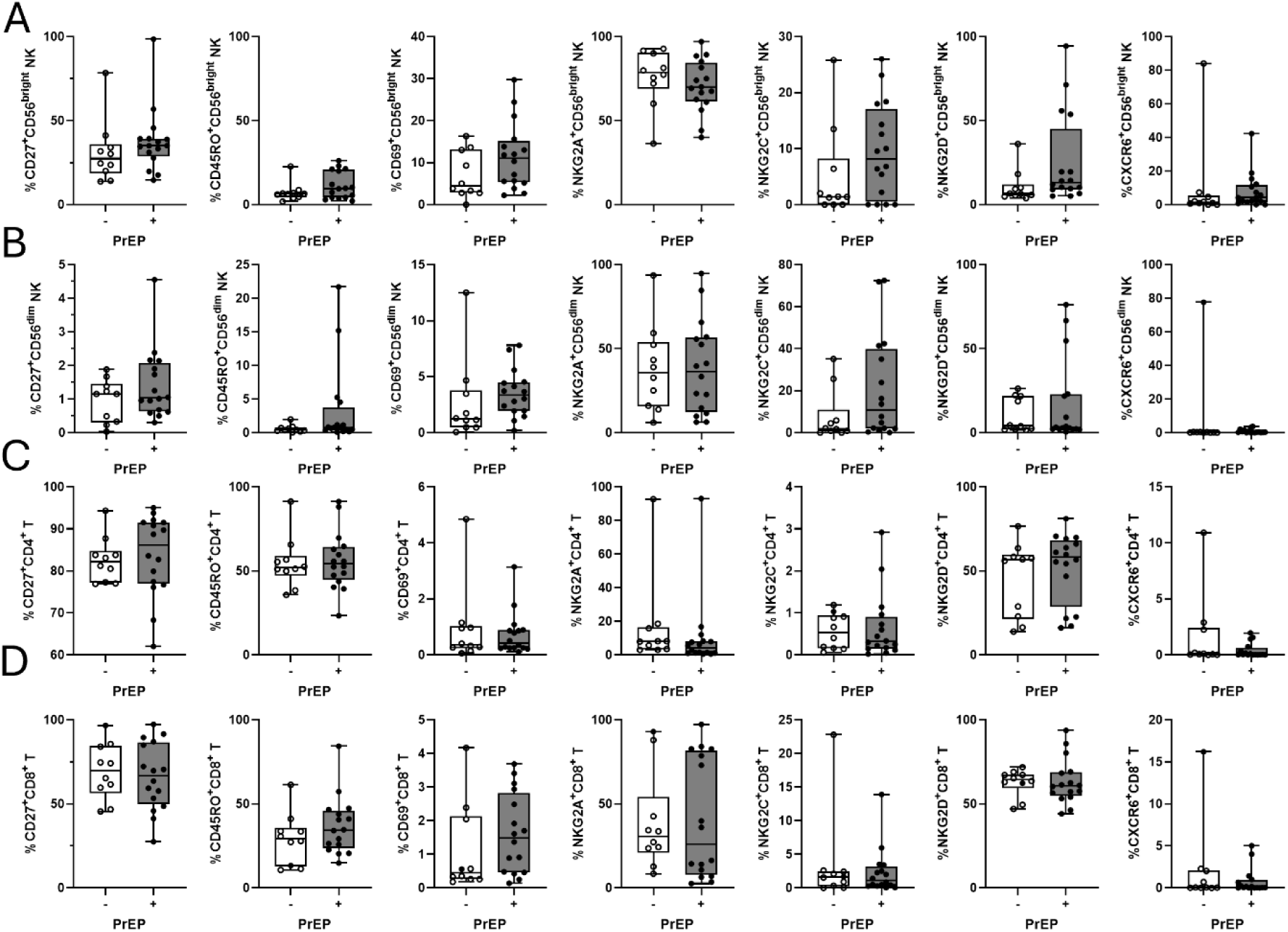
Daily PrEP does not alter lymphocyte phenotypic profiles. PBMC were isolated from peripheral blood of PrEP naive individuals (white; n=11) or individuals established on PrEP (grey; n=15) and stained with fluorochrome conjugated antibodies for CD56, CD3, CD4, CD8, CD27, CD45RO, CD69, NKG2C, NKG2D, CXCR6, for flow cytometric analysis. The frequency of **(A)** CD56^bright^ NK cells, **(B)** CD56^bright^ NK cells, **(C)** CD4^+^ T cells, or **(D)** CD8^+^ T cells expressing CD27, CD45RO, CD69, NKG2C, NKG2D or CXCR6. Each dot represents an individual donor, and data were analysed using an unpaired t test; however, results were not statistically significant.

**Supplemental Figure 2.**
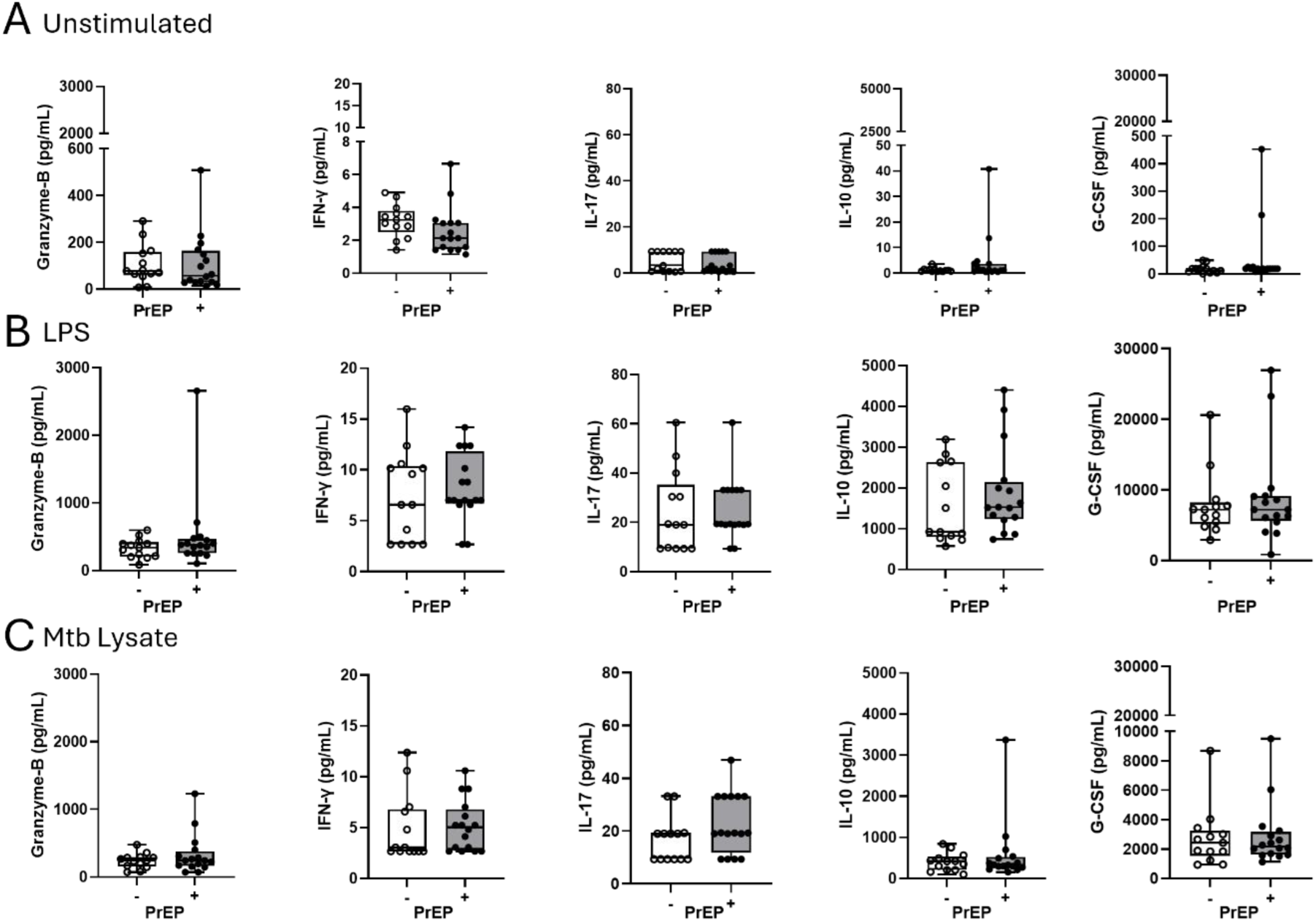
Monocyte functional outputs that remain unchanged in individuals established on PrEP. Monocytes were magnetically sorted from the PBMC of PrEP naive individuals (white; n=11) or individuals established on PrEP (grey; n=15) and **(A)** left unstimulated, **(B)** stimulated with LPS or **(C)** stimulated with Mtb lysate for 18 hours. The concentrations (pg/ml) of Granzyme-B, IFN-γ, IL-17, IL-10 or G-CSF in the supernatant were measured using a Luminex Discovery assay. Each dot represents an individual donor, and data were analysed using an unpaired t test; however, results were not statistically significant.

**Supplemental Figure 3.**
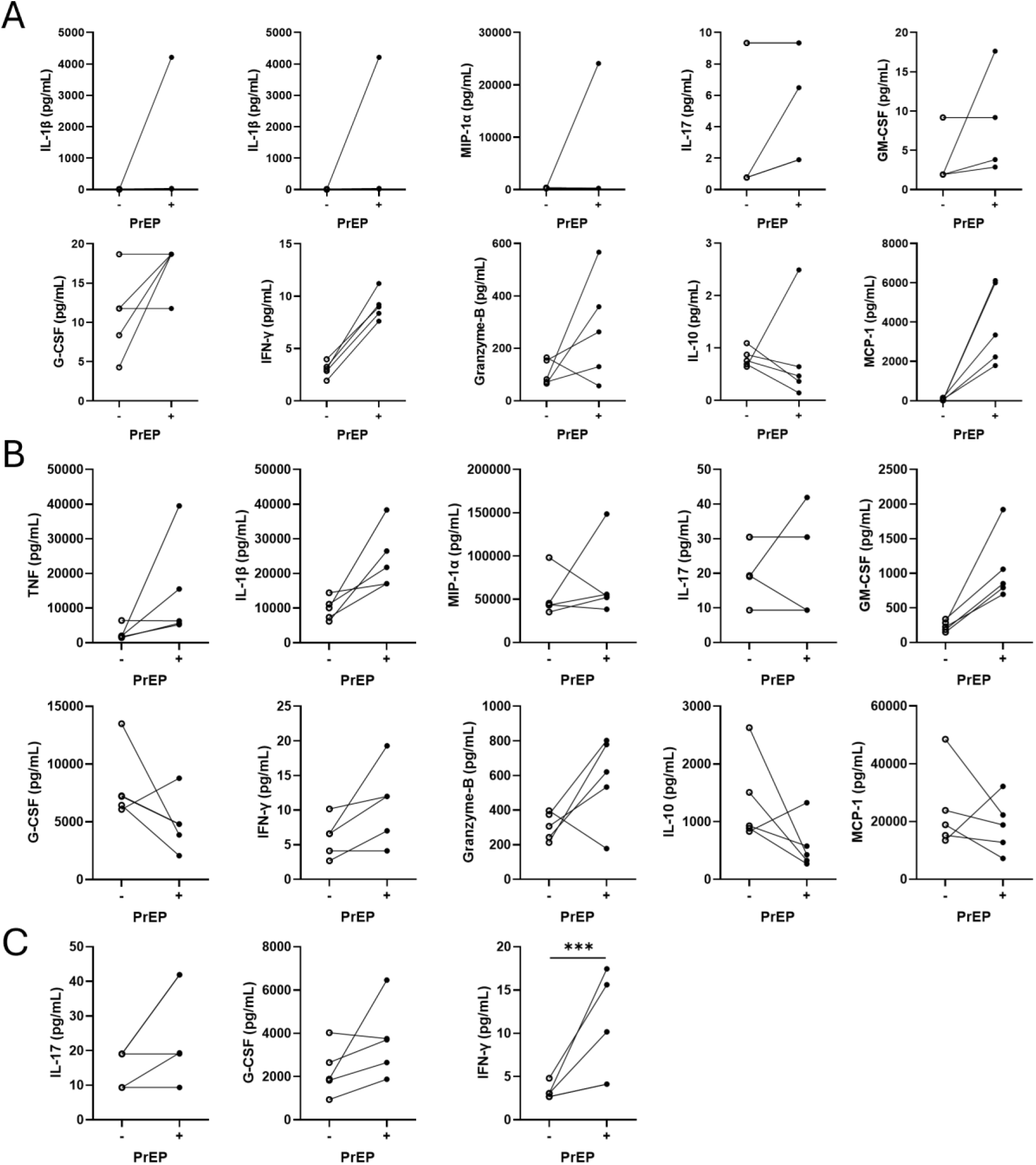
The effect of PrEP on monocyte functional output in response to diverse stimuli. PBMC were isolated from the matched blood (n=5) of PrEP naïve individuals at T1 (-; open circle) who were established on PrEP at T2 (+; closed circle). Magnetically sorted monocytes were **(A)** left unstimulated, **(B)** stimulated with LPS or **(C)** stimulated with Mtb lysate for 18 hours. The concentrations (pg/ml) of TNF, IL-1β, MIP-1α, IL-17, GM-CSF, G-CSF, IFN-γ, Granzyme-B, IL-10, and MCP-1 in the supernatant were measured using a Luminex Discovery assay. Each dot represents matched data at each timepoint joined by a line. Data were analysed using an unpaired t test where *P<0.05, **P<0.01, or ***P<0.001.

## Notes

### Competing Interest Statement

The authors have declared no competing interest.

